# Apicomplexans predict thermal stress mortality in the Mediterranean coral *Paramuricea clavata*

**DOI:** 10.1101/2022.11.23.517658

**Authors:** Anthony M Bonacolta, Jordi Miravall, Daniel Gómez-Gras, Jean-Baptiste Ledoux, Paula López-Sendino, Joaquim Garrabou, Ramon Massana, Javier del Campo

**Affiliations:** Department of Marine Biology and Ecology, Rosenstiel School of Marine, Atmospheric and Earth Science, University of Miami, Miami, Florida, 33149 USA; Institut de Biologia Evolutiva (CSIC-Universitat Pompeu Fabra), Barcelona, Catalonia, Spain; Institut de Ciències del Mar – CSIC, Barcelona, Catalonia, Spain; Hawai‘i Institute of Marine Biology, University of Hawai‘i at Mānoa, Kaneohe, HI, USA; Departament de Biologia Evolutiva, Ecologia i Ciències Ambientals, Institut de Recerca de la Biodiversitat (IRBIO), Universitat de Barcelona, Barcelona, Spain; CIIMAR/CIMAR, Centro Interdisciplinar de Investigação Marinha e Ambiental, Universidade do Porto, Porto, Portugal.

**Keywords:** Microbiome, 18S rRNA, Metabarcoding, Octocorallia, *P. clavata*, Apicomplexa, Protists, Syndiniales, Cnidaria, Coral

## Abstract

The octocoral *Paramuricea clavata* is an ecosystem architect of the Mediterranean temperate reefs that is currently threatened by episodic mass mortality events related to global warming. Local average thermal regimes nor recent thermal history have been shown to play a significant role in population thermotolerance in this species. The microbiome, however, may play an active role in the thermal stress susceptibility of corals, potentially holding the answer as to why corals show differential sensitivity to heat-stress. To investigate this, the prokaryotic and eukaryotic microbiome of *P. clavata* collected from around the Mediterranean was characterized before experimental heat-stress to determine if its microbial composition influences the thermal response of the holobiont. We found that the prokaryotic community was not informative in predicting the thermal susceptibility of *P. clavata*. On the other hand, members of *P. clavata’s* microeukaryotic community were significantly correlated with thermal stress sensitivity. Syndiniales from the *Dino-Group I Clade 1* were significantly enriched in thermally resistant corals, while the apicomplexan corallicolids were significantly enriched in thermally susceptible corals. Corallicolids are associated with 70% of coral genera around the world, yet the ecological role of this general anthozoan symbiont has yet to be determined. We hypothesize that *P. clavata* mortality following heat-stress may be caused by a shift from apparent commensalism to parasitism in the corallicolid-coral host relationship driven by the added stress. Our results show the potential importance of corallicolids and the rest of the microeukaryotic community of corals to understanding thermal stress response in corals and provides a useful tool to guide conservation efforts and future research into coral-associated microeukaryotes.

## Background

*Paramuricea clavata* is a slow-growing octocoral prevalent in the temperate Mediterranean Sea [1]. Gorgonians, such as *P. clavata*, are habitat-forming species of biodiversity-rich hard bottom benthic communities where they provide complex habitat for numerous organisms contributing significantly to the biodiversity of the region [2]. *P. clavata’s* slow growth rate, long population turn-over time, high post-settlement mortality, and low genetic connectivity puts it at high-risk of extinction caused by natural and anthropogenic stressors [1]. *P. clavata* populations have been facing increased mortality due to episodic mass mortality events (MMEs) [3]. MMEs, which were once uncommon in temperate ecosystems such as the Mediterranean Sea, have been increasing since the turn of the century, with most of these occurring in the Western Mediterranean [4]. The increase in the frequency, duration, and intensity of marine heatwaves driven by global warming have been suggested to be the primary cause of the increase in MMEs in the Mediterranean. Thus, MMEs are leading to the current biodiversity crisis in the region [5, 6]. *P. clavata* are heavily affected by MMEs with mortality sometimes reaching 80% in affected populations and recovery hindered by the recurrence of marine heatwaves. The loss of this habitat-forming species causes a functional shift in Mediterranean hard-bottom communities leading to simplified habitats with lower species diversity [7, 8]. To prevent the significant decline of *P. clavata* and other habitat forming octocorals in the Mediterranean, the effect of thermal stress on the *P. clavata* holobiont must be better understood.

Thermal stress typically causes death in anthozoans by disrupting metabolic pathways, increasing physiological stress, decreasing host defense, causing energy constraints, and promoting the growth of opportunistic pathogens [9–11]. In *P. clavata,* exposure to higher temperatures causes usually partial or full necrosis of colonies, where branches first change color from red to grey before denudation of the skeleton [12]. This contrasts with “bleaching” which takes place in zooxanthellate corals and is when the algal symbiont leaves the coral host during periods of high heat-stress [13]. *P. clavata* is more sensitive to elevated seawater temperatures than other gorgonians, marked by a significant loss in energy reserves, reduced feeding ability, and partial mortality after heat-stress [14]. Furthermore, a companion study using the same *P. clavata* from this research found no significant association between local average thermal regimes nor recent thermal history and population thermotolerance; however, inter-individual variation within populations was found [15]. A complete characterization of the *P. clavata* microbiome before heat-stress can better elucidate the role of the microbiome to overall holobiont health and fitness in the face of warming ocean temperatures.

Some octocoral-associated microorganisms are considered beneficial members of the coral holobiont contributing to nitrogen fixation, sulfur-cycling, antibiotic production, and host defense [16]. *P. clavata* has a consistent composition of the bacterial component of the microbiome across regions, dominated (above ∼90% relative abundance) by the common coral symbiont *Endozoicomonas,* a gammaproteobacterium of the order Oceanospirillales [17, 18]. Notably, *P. clavata*’s bacterial community differs significantly from the microbiome of the water surrounding the organism and that of neighboring octocorals, such as *Eunicella* [19]. Other bacteria commonly found within *P. clavata* include Alphaproteobacteria, Betaproteobacteria, and Actinobacteria [17]. Despite the research into the bacterial community found in *P. clavata* throughout the Mediterranean Sea, there has been no research into the microeukaryotic community of this coral which constitutes a significant portion of the microbial diversity in other corals and may contribute significantly to holobiont fitness in a changing climate [20].

Microeukaryotes (protists and unicellular fungi) are important components of many of the Earth’s ecosystems, including being the main contributors to oceanic primary production and carbon mobilization in the microbial loop [21]. They may also play an important role in many marine holobionts. The corallicolids, which are part of a lineage of parasitic alveolates known as the Apicomplexa are associated with 70% of screened coral genera globally [22]. Apicomplexans also include such well-known human parasites like *Plasmodium*, the causative agent of malaria, and *Toxoplasma*, a wide-spread mammal-infecting pathogen. While the corallicolids, based on their distribution and presence in healthy individuals [23], seem to be commensals in the coral microbiome, their evolutionary context suggests a more parasitic role for this general anthozoan associate.

To better understand the microbiome of *P. clavata* and its ability to predict thermal stress susceptibility we conducted a common garden heat-stress experiment using corals collected from 12 populations (148 total individuals) across a wide geographic range of *P. clavata* (See Supplementary Figure 1, Additional File 1). We used an 18S rRNA and 16S rRNA gene metabarcoding approach to get a holistic view of the microbes which may confer thermal resiliency to *P. clavata*.

## Methods

### Sample Collection

Branch fragments (30 colonies per population) of healthy adult *P. clavata* were collected from four localities (11 discrete populations) by scuba across the 18^th^-19^th^ of September 2017 encompassing much of the known Mediterranean distribution of the octocoral. From Croatia, *P. clavata* populations from Balun (Lat, Lon; 43.803889, 15.255000) and Mana (43.800273, 15.266389) were sampled. Three populations were sampled from Portofino (Liguria, Italy): Altare (44.308918, 9.178645), Indiano (44.312476, 9.166889), and Lighthouse (44.298650, 9.219014). Three populations were sampled from Scandola (Corsica, France): Palazzinu (42.379854, 8.549999), Gargallu (42.371839, 8.534394), and Palazzu (42.379873, 8.546163). Three populations were sampled from Medes Islands (Catalonia, Spain): La Vaca (42.048047, 3.226322), Pota de Llop (42.0497, 3.2254), and Tascons (42.042189, 3.2269). Lastly, a population thought to be *P. clavata* was sampled from an Atlantic locality from Sagres, Portugal (37.012129, -8.924541). While underwater, samples were initially stashed in small plastic jars or bags (no more than 10 per container) until being brought to the surface. Characteristic information from each site can be found in Supplementary Table 1 (See Supplementary Table 1, Additional File 2). Sampling was conducted as described in Gómez-Gras et al. 2022 [15]. In short, 0.5 cm branch fragments were sampled for microbiome analyses at time of collection, stored in a 2 mL cryovial containing 96% EtOH, flash-frozen using liquid nitrogen, and shipped to the Institut de Ciències del Mar (CSIC) in Barcelona, Catalonia, Spain on ice. 10-17 individuals per population were included in microbiome analyses, depending on their availability. 10 cm long coral specimens were maintained in a recirculating aquaria system overnight, then transferred to 2 L bags containing water (10 coral fragments per bag). Bags were placed into polystyrene boxes (3 bags per box) then shipped to Barcelona. For the coral specimens from Medes and Scandola, the coral were transported in coolers immediately after sampling, skipping the overnight aquaria step of the other populations. Transportation times between the sampling location and CSIC ranged from several hours (Medes) to 36 hours (Portofino).

### Thermal Stress Experiment

The experiment was conducted in the Aquarium Experimental Zone of the Institut de Ciències del Mar in Barcelona (ICM-CSIC). Sets of 70 L aquarium tanks were divided between control and treatment groupings, each containing 3 replicate tanks per population with 10 coral fragments in each (30 individuals in total per population and experimental condition). To establish and maintain experimental conditions, each tank has individual heaters, temperature controllers, water pumps, and HOBO temperature data logs (recorded every 5 mins). Coral fragments were first acclimated to the aquarium environment for 2 months before experimentation began. Any coral showing signs of disease or necrosis during this period was promptly discarded before experimentation began. For the control tanks, water temperature was maintained at 16-18°C for the entirety of the experiment. For the treatment tanks, the temperature increased step-wise from 18°C to 25°C over the course of three days. Water for both treatments was sourced from a buffer tank with 50 mm sand-filtered Mediterranean seawater pumped from 15 m depth. The coral fragments were fed three times per week until the end of the experiment. Feeding alternated between 3 mL of Bentos Nutrition Marine Active Supplement (Maim, Vic, Spain) and a tablet of frozen copepods (*Cyclops* spp.; Ocean nutrition, Antwerp, Belgium). Following 21 days of experimentation, the responses of individual coral fragments were assessed by measuring the percentage of tissue necrosis. Percent necrosis per fragment was assessed on days 14 and 21. Thermal response for each individual was categorized as follows by interpretation of the results of the two necrosis measurements: Super Resistant (0% necrosis), Resistant (< 25% necrosis), Tolerant (25-75% necrosis), Sensitive (>75% necrosis), and Hyper Sensitive (100% necrosis). Supplementary Table 1 (See Supplementary Table 1, Additional File 3) shows the thermal resiliency categories for each coral sample at the end of the experiment.

### DNA Extraction and rRNA Gene Metabarcoding

*P. clavata* microbiome samples were thawed at room temperature and then processed using a modified DNeasy Mini Kit (QIAGEN) protocol. In short, samples were digested for 3 hours at 56°C with slow agitation in 180 µl of tissue lysis buffer and 20 µl of proteinase K. During this time samples were mixed frequently by vortexing. Once the incubation ended 200 µl of Lysis buffer was added to the tissue lysates followed by 200 µl of 96-100% ethanol. Tissue lysates were then mixed by vortexing and the resulting mixtures were pipetted (discarding the central axis and the sclerites released from the tissues) into DNeasy Mini spin columns placed in 2 ml collection tubes. The columns were centrifuged for 1 minute at 8000 rpm and room temperature. The collection tubes with the mixture were discarded and the columns were placed in new collection tubes. The samples contained in the DNeasy Mini spin columns were then washed in two steps. First, 500 µl of Wash buffer 1 was added to the columns followed by a centrifugation of 1 minute at 8000 rpm and room temperature. Second, 500 µl of Wash buffer 2 was added and columns were centrifuged for 3 minutes at 14000 rpm and room temperature. Finally, to elute the DNA the columns were placed in 1.5ml Eppendorf tubes and 200 µl of Elution buffer was added to the center of the spin column membrane. Lastly, the columns were incubated for 1 minute at room temperature and then they were centrifuged for 1 minute at 8000 rpm at room temperature. DNA concentration from purified samples was measured using Qubit fluorometer. Concentrations that were higher than 1 ng/µl were considered acceptable for PCR. The V4 region of the prokaryotic 16S rRNA gene was amplified according to the Earth Microbiome Project protocol [53] and the V4 region of the eukaryotic 18S rRNA gene was amplified using a nested PCR approach validated in del Campo et al. 2019 [54]. Briefly, for the 18S rRNA metabarcoding, the samples were first amplified with UNonMet-PCR primers which reduce the metazoan signal (18S-EUK581-F 5’-GTGCCAGCAGCCGCG-3′ 18S-EUK1134-R 5’-TTTAAGTTTCAGCCTTGCG-3′), then reamplified with primers from Comeau et al. 2011 [55]. PCR mixture for one sample contained 4 µl 5X HS Buffer, 0.4 µl dNTP, 1 µl each of the forward and reverse primers (diluted at 1:10), 0.6 µl DMSO, 0.2 µl Phusion DNA Polymerase, 3 µl of the sample DNA template, and reaction volumes made up to 20 µl with MilliQ water (9.8 µl). The PCR program of the thermal cycler was set up as follows: an initial denaturation step at 98°C for 30 seconds, subsequent 35 cycles of denaturation at 98°C for 10 seconds, annealing at 51.1°C for 30 seconds and elongation at 72°C for 1 minute, all of them followed by a final elongation at 72°C for 5 minutes. Gel electrophoresis was used to ensure PCR amplification was successful. Electrophoresis gel was made with 1g of ultrapure agarose dissolved in 100 ml of 1X Tris-acetate-EDTA buffer (TAE) and stained with 5 µl of SYBR safe DNA gel stain. The gelling process has been carried out at room temperature for about 20 minutes and the agarose gel was ran at 100 V for about 20 minutes at room temperature. The agarose gel was finally visualized under UV light using a Bio-Rad Chemi XRS Gel Documentation system and Bio-Rad Quantity One software. Those samples that didn’t show a clear band on the agarose gel were amplified again adding 0.2 µl MgCl2 50mM to the PCR mixture for one sample. The rest of the components of the PCR mixture for one sample were maintained at the same volumes as before and reaction volumes made up to 20 µl with MilliQ water (9.6 µl). The PCR program of the thermal cycler was set up as before and PCR samples were checked again by an electrophoresis. The PCR products were cleaned using QIAquick PCR Purification Kit (Qiagen). The buffer PB of this purification kit was used with a pH indicator added to it. A volume of 8 µl of the PCR sample was mixed with 40 µl of the Buffer PB and transferred to a QIAquick column. Columns were centrifuged for 60 s to bind DNA to the column membrane. Next columns were washed by adding 750 µl of Buffer PE and centrifuging them in two steps of 60 s each one. Finally, DNA was eluted by adding 50 μ of Buffer PB and centrifuging the columns for 1 minute. DNA concentrations from purified PCR samples were checked using Qubit fluorometer before being sent to the Integrated Microbiome Resource facility at the Centre for Comparative Genomics and Evolutionary Bioinformatics at Dalhousie University for further 18S rRNA amplification and sequencing. 96 holes microplates were prepared in order to send them to Dalhousie University for sequencing on a Illumina MiSeq using 300+300 bp paired-end V3 chemistry. Plates were prepared following the receiver instructions. Plate holes were filled with 10 μ DNA purified samples and the PCR purified samples and plates were sealed using an adhesive plastic seal.

### Microbiome Analysis

Raw sequencing data for this project has been deposited on NCBI (BioProject: PRJNA928446). Primers were removed from reads using Cutadapt v3.1 [56]. The trimmed reads were then processed in R using DADA2 [24]. For the 18S rRNA gene amplicons, forward reads were truncated at 230 bp and reverse reads were truncated at 195 bp. Reads were then denoised and joined into amplicon sequence variants (ASVs). Chimeras were removed using the ‘removeBimeraDenovo’ command with the “consensus” option. Taxonomy was assigned to 18S ASVs in DADA2 using PR2 v4.13.0 [57]. For the 16S rRNA gene amplicons, forward reads were truncated at 250 bp and reverse reads at 210 bp. Truncated reads were then denoised and joined into ASVs before assigning taxonomy using SILVA v138 [58] in DADA2. Sequence tables, taxonomy tables, and metadata for both the 16S and 18S rRNA gene amplicon datasets were uploaded into a Phyloseq object in R for further filtering and analysis [59]. For the 16S rRNA gene amplicons, ASVs corresponding to Chloroplast and Mitochondria were removed. One sample was also removed for having under 250 reads. Additionally, ASVs were kept in the dataset only if they were in the most abundant 99% of taxa in at least one of the samples. Following this filtering 4 546 ASVs and 92 samples remained. The 18S amplicon were subjected to the same filtering parameters except ASVs corresponding to Metazoa and Embryophyceae were also removed from the dataset, resulting in 3181 ASVs across 138 samples. Bubble plots, alpha-diversity, and beta-diversity figures were constructed using ggplot2 and tidyverse packages [60]. Statistical analyses were conducted using the microbiomeMarker package and vegan [61, 62]. Analysis of similarities (ANOSIM) within vegan was used to test for differences in beta-diversity between groups using 999 permutations. Linear Discriminant Analysis Effect Size (LEfSe) [63] and Analysis of Compositions of Microbiomes with Bias Correction (ANCOM-BC) [64] were used to find significantly enriched and differentially abundant taxa between treatment groups. For the ANCOM-BC test, region of origin was treated as a fixed variable and the p-values were adjusted using false discover rate (“fdr”).

In order to place the corallicolid ASVs in a phylogenetic context, a multiple sequence alignment (MSA) was created using publicly available full-length 18S rRNA sequences of corallicolids and the program MAFFT. Abundant corallicolid ASVs (more than 50% of corallicolid signal in at least one sample) were then extracted from the dataset. Hmmalign was then used to map these ASVs to the full length 18S rRNA corallicolid MSA [65]. A maximum likelihood phylogenentic tree was constructed with RAxML using this MSA (GTRCAT substitution model, N = 1 000) [66]. The resulting tree was modified for publication using ggtree [67]. For the dinoflagellate and ciliate trees, a similar approach was taken, except no publicly available sequences were used in their construction and alignment.

To assess cross-domain positive-correlation in the occurrence patterns of ASVs, the 16S and 18S rRNA gene amplicon tables were combined for samples present in both datasets. After setting a filter prevalence threshold of 0.001, correlation patterns were inferred for the combined dataset using SPARCC which considers the compositional nature of metabarcoding data [68]. A network was then constructed using SpiecEasi with a p-value threshold of 0.05 (ref. 30). The top families exhibiting positive co-occurrence relationships were plotted in a chordplot using the MicroEco package [70].

## RESULTS

### The bacterial microbiome of *Paramuricea clavata*

For the 16S rRNA gene sequencing, 1332678 total reads were recovered after processing in DADA2 [24]. Following filtering, a total of 4540 Amplicon Sequence Variants (ASVs) across 92 samples were left for bacterial community analysis. The water and food microbiome samples were taxonomically distinct from the coral microbiome samples in this study (See Supplementary Figure 2, Additional File 4). The population originally sampled from Sagres, Portugal exhibited a distinct microbiome from the other *P. clavata* populations (Fig. 1) and was later determined to be a different species based on genomic work [25]. It was hence removed from further analysis. Oceanospirillales, mostly consisting of *Endozoicomonas*, constituted most of the bacterial community of *P. clavata* (Fig. 1). Other highly abundant bacterial orders include Alteromonadales, Rhodobacterales, and Flavobacteriales. A high number of unidentified bacterial ASVs were also present in the dataset. Subsequent BLAST analysis of these ASVs yielded no definitive taxonomic identification.

**Figure 1.**
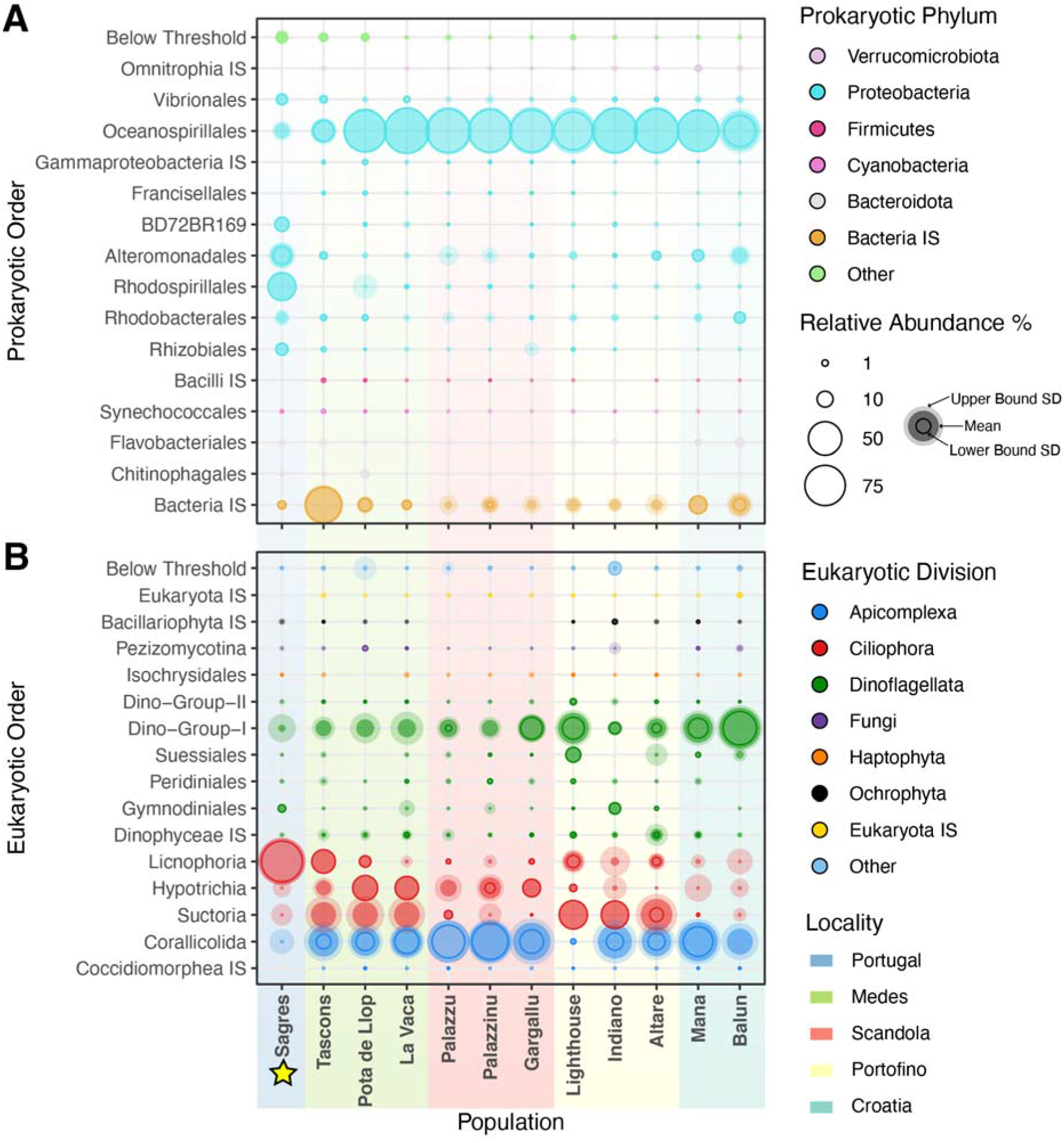
Relative abundance of microbial orders within *P. clavata* across its geographic distribution. According to 16S rRNA gene amplicons (A) and 18S rRNA gene amplicons (B) metabarcoding. *P. clavata* populations are listed from west to east, with major localities color coded. Sagres is denoted with a star as this population was excluded from further analysis since it is a different species. Microbial taxa are ordered taxonomically. Only those taxa with a median relative abundance above 1% are shown, the rest are listed as “other”. IS is short for Incertae sedis, meaning there is uncertainty in the taxonomic position within the broader taxonomic group. Relative abundance bubbles are shown at 80% opacity and correspond to within domain (Prokaryotic or Eukaryotic) relative abundance. The upper-bound of the standard deviation is shown by the opaquer outer bubble. The lower-bound of the standard deviation is shown by a solid circle. Lower bounds with a negative value are not shown.

Pair-wise analysis of the bacterial communities at the ASV-level between localities showed significant differences in Shannon-Wiener alpha-diversity (Fig. 2). Additionally, the beta-diversity of the bacterial community between localities was significantly different as well (ANOSIM; P = 0.001; Fig. 3). Overall, the bacterial community of *P. clavata* clustered closely based on geographic origin.

**Figure 2.**
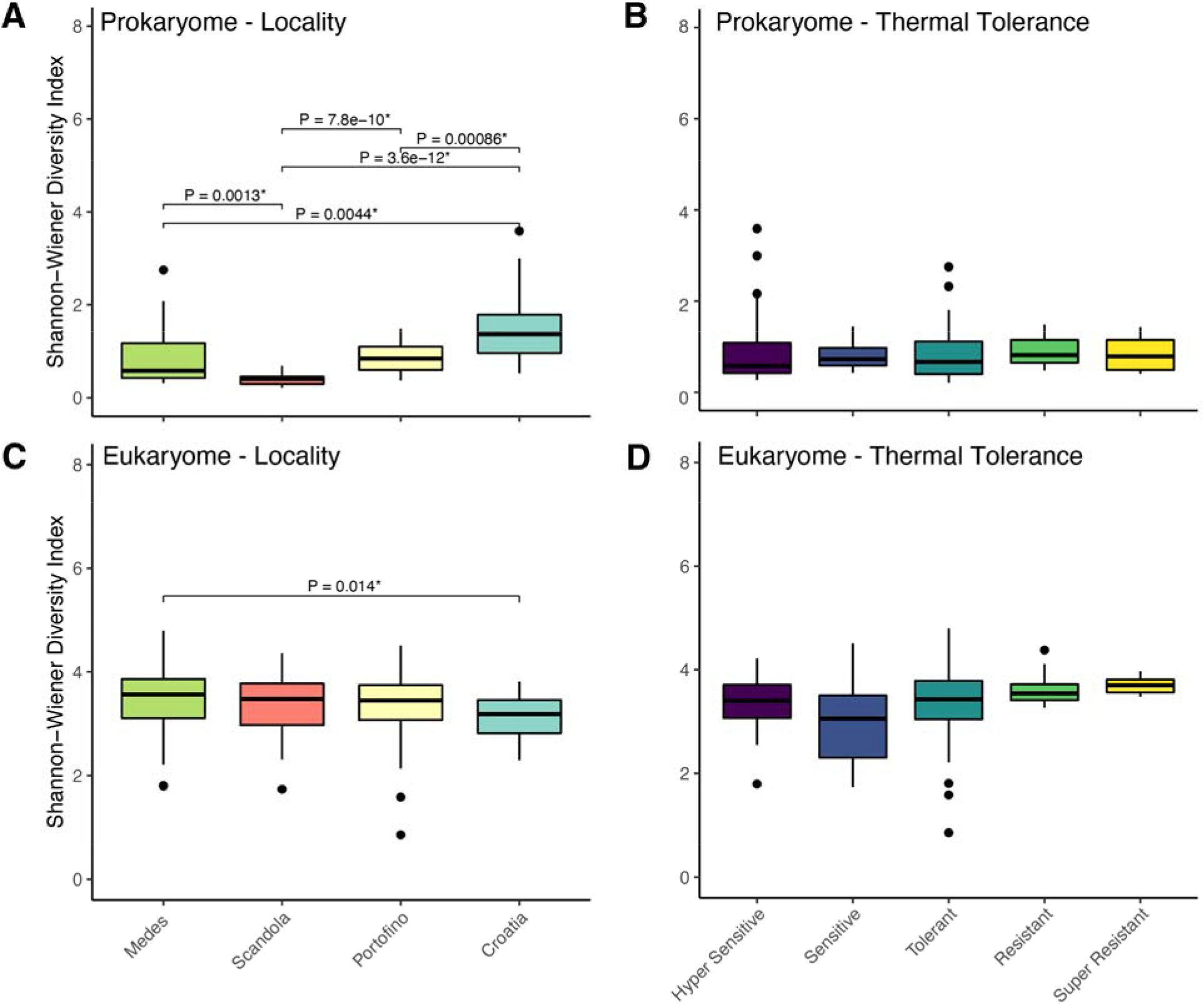
Shannon-Wiener Alpha Diversity of the prokaryome and eukaryome of *P. clavata* across sampled localities and thermal tolerance categories. Mean comparison p-values are added to significant group comparisons.

**Figure 3.**
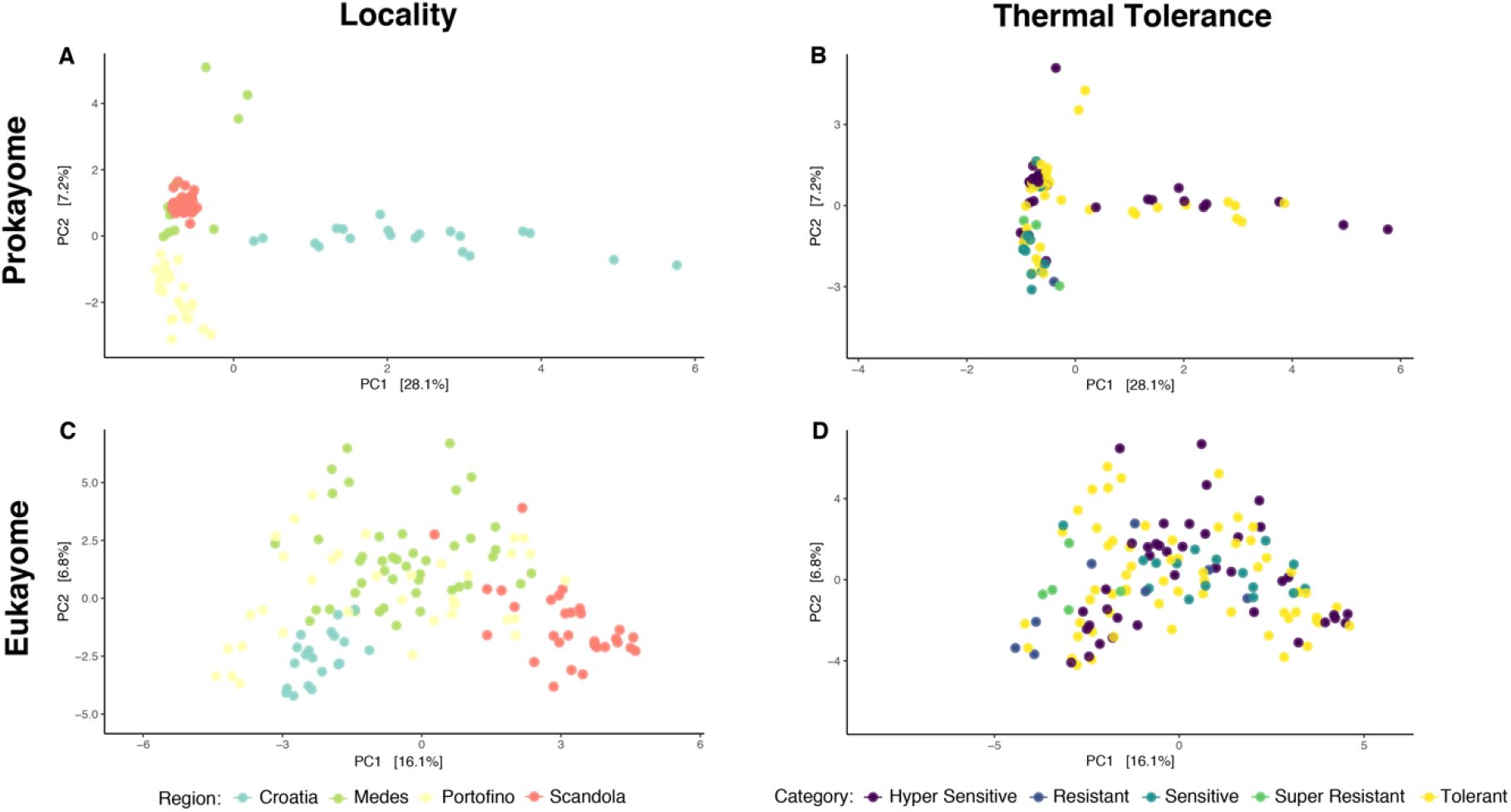
Aitchison distance PCAs of the prokaryome and eukaryome of *P. clavata* across sampled localities and thermal tolerance categories. Analysis of similarities (ANOSIM) within vegan was used to test for differences in beta-diversity between groups using 999 permutations. The beta-diversities of both the bacterial and microeukaryotic communities between localities were significant (ANOSIM; P = 0.001).

The dominance of *Endozoicomonas* (Proteobacteria; Oceanospirillales) in the bacterial community of *P. clavata* masked any fine-scale differences in the bacterial community composition across the range of thermal susceptibilities (Fig. 4). Analyzing the bacterial community with *Endozoicomonas* removed, did not reveal any noticeable pattern in taxon composition in relation to thermal stress susceptibility (See Supplementary Figure 3, Additional File 5). Additionally, an ANCOM-BC analysis of the Prokaryotic Orders corrected for the region of origin and false discovery rate did not identify any significant bacterial taxa between the thermal susceptibility categories.

**Figure 4.**
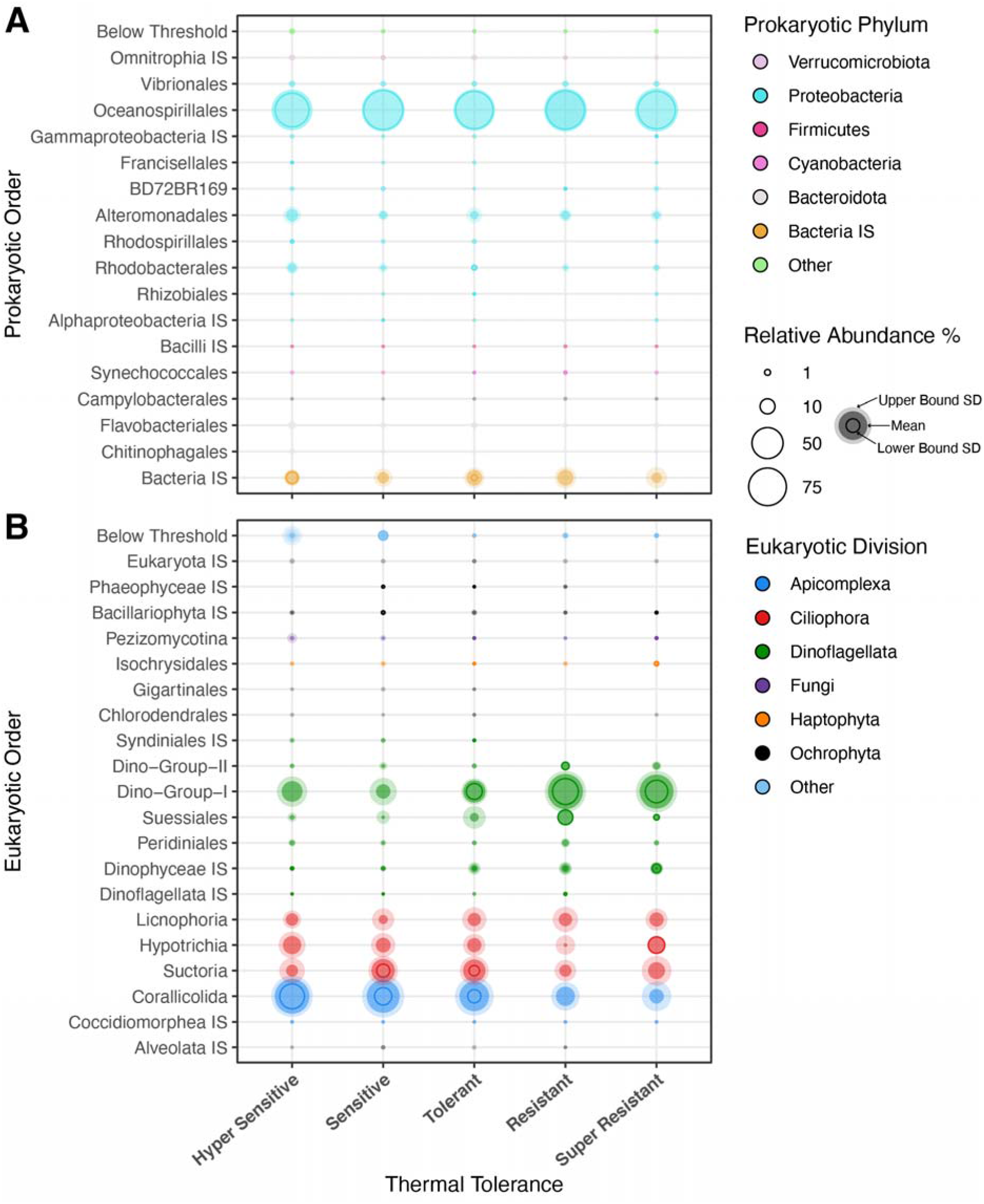
Relative abundance of microbial orders within *P. clavata* across thermal susceptibility categories. According to 16S rRNA gene amplicons (A) and 18S rRNA gene amplicons (B) metabarcoding. The Sagres population was not included in this analysis (see Methods section). Hyper Sensitive (n_bac_ = 29; n_euk_ = 39) corals were those which showed 100% necrosis during the heat experiment. Sensitive (n_bac_ = 3; n_euk_ = 16) corals were those that showed < 75% necrosis. Tolerant (n_bac_ = 42; n_euk_ = 61) corals were those that showed 25-75% necrosis. Resistant (n_bac_ = 3; n_euk_ = 7) corals showed < 25% necrosis at the end of heat-stress. Super Resistant (n_bac_ = 4; n_euk_ = 5) corals showed 0% necrosis at the end of heat-stress. Microbial taxa are ordered taxonomically. Only those taxa with a median relative abundance above 1% are shown, the rest are listed as “other”. IS is short for Incertae sedis, meaning there is uncertainty in the taxonomic position within the broader taxonomic group. Relative abundance bubbles are shown at 80% opacity and correspond to within domain relative abundance. The upper-bound of the standard deviation is shown by the opaquer outer bubble. The lower-bound of the standard deviation is shown by a solid circle. Lower bounds with a negative value are not shown.

### The eukaryotic microbiome of *Paramuricea clavata*

For the 18S rRNA sequencing, 18955358 total reads were recovered, resulting in 3181 ASVs across 138 samples following filtering. This community is dominated by 3 major clades of Alveolates: Apicomplexans, Ciliates, and Syndiniales (Fig. 1). One of the primary microeukaryotic taxa present, Dino-Group-I (Dinoflagellata; Syndiniales), exhibited its highest relative abundance in the more Eastern populations (Balun, Mana, & Lighthouse) (Fig. 1). Corallicolids (Apicomplexa; Corallicolida) exhibited a high relative abundance across every population sampled, excluding Lighthouse (Fig. 1). Additionally, three distinct genera of ciliates (Ciliophora) were found across the surveyed geographic range of *P. clavata*, with *Licnophoria* spp. being prevalent at Tascons, Lighthouse, and Altare (Fig. 1). *Hypotrichia* spp. were found mainly in the Western populations (Tascons, Pota de Llop, La Vaca, Palazzu, Palazzinu, & Gargallu) (Fig. 1). Lastly, *Suctoria* spp. were most prevalent within the *P. clavata* of the Medes and Portofino localities (Fig. 1).

At the ASV-level, the locality-level microeukaryotic communities of *P. clavata* exhibited higher Shannon-Wiener alpha-diversity compared to the bacterial communities (Fig. 2). Additionally, the microeukaryotic community only showed a significant difference in the alpha-diversity of *P. clavata* between Croatia vs Medes. Furthermore, the beta-diversity between localities was significantly different for the microeukaryotic community (ANOSIM; P = 0.001; Fig. 3).

Syndiniales were enriched in the super-resistant corals (LEfSe; P = 0.0016). Specifically, *Dino-Group I Clade 1* abundance in the holobiont was associated with the increased thermotolerance in *P. clavata* and was enriched in the super-resistant individuals (LEfSe; P = 0.0027; Fig. 4). At the ASV level, ASV_20 and ASV_13 were the *Dino-Group I Clade 1* ASVs which were highly abundant in the resistant groups relative to the other Alveolates (See Supplementary Figure 4, Additional File 6). Concurrently, a high relative abundance of corallicolids within the holobiont was associated with higher sensitivity to thermal stress (Fig. 4). LEfSe analysis revealed that the *Corallicolidae* family was significantly enriched in the sensitive corals (LEfSe; P = 0.015). ASV_6, within the COR3 clades of corallicolids, showed the highest abundance amongst of the alveolates in the hyper-sensitive corals (See Supplementary Figure 4, Additional File 6). ANCOM-BC analysis of the microeukaryotic Orders corrected for the region of origin and false discovery rate also confirmed these associations, with corallicolids being significantly more abundant (∼2.3 natural log fold higher abundance; ANCOMBC, p_adj_ = 0.0003) in the sensitive and hypersensitive corals compared to the resistant and hyper resistant ones, while Dino-Groups I and II showed an opposite pattern (Fig. 5; ∼1.4 natural log fold lower abundance; ANCOMBC, p_adj_ = 0.04). On the other hand, ciliates showed a steady abundance throughout thermal sensitivity groups, except for ASV_7 (*Ephelota mammillata*) which was highly abundant in the sensitive and tolerant groupings (See Supplementary Figure 4, Additional File 6). Alpha-diversity and beta-diversity analysis of both the bacterial community and microeukaryotic community yielded no significant differences between any of the thermal tolerance categories (Figs. 2 & 3).

**Figure 5.**
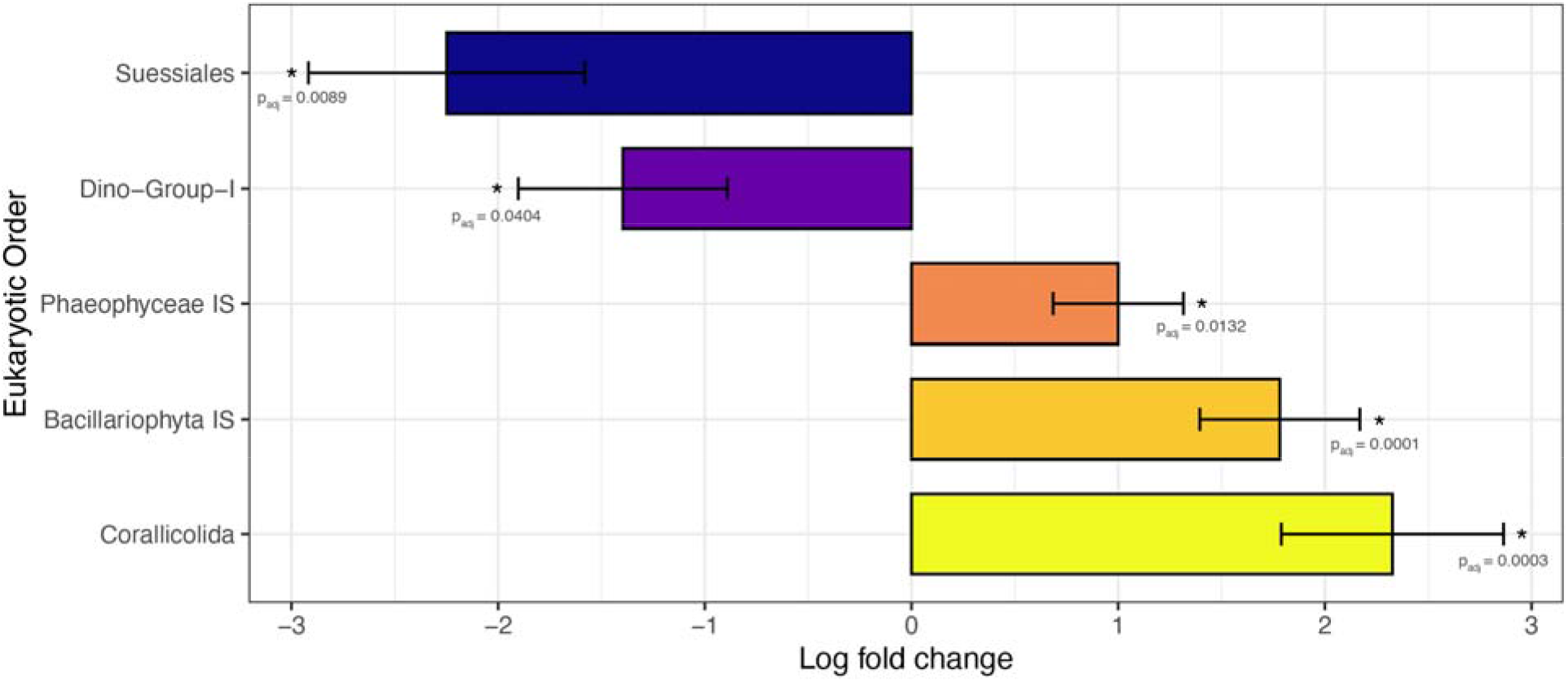
ANCOMBC Significant Taxa. Significant differentially abundant microbial taxa between sensitive (hyper sensitive + sensitive; n_euk_ = 55) vs resistant (super resistant + resistant; n_euk_ = 12) samples according to ANCOMBC analysis while controlling for region of origin and false discovery rate. Natural log fold differences in abundance for each significant taxa are shown with SD. Adjusted p-values are shown. IS is short for Incertae sedis, meaning there is uncertainty in the taxonomic position within the broader taxonomic group.

### Phylogenetic placement of the microeukaryotes

Phylogenetic analysis of the most prevalent Alveolate ASVs found within *P. clavata* revealed a diversity of clades associated with the octocoral across the surveyed geographic range. ASV_6, which was the most abundant corallicolid ASV and found in most samples, was most abundant in the Scandola populations (Fig. 6). An evolutionary placement algorithm based on RAxML placed this ASV as closely related to MN022452_Corallicolidae_COR3_sp. (99.22% Similarity), a sequence recovered from the deep-sea octocoral, *Callogorgia delta* in the Gulf of Mexico. A second clade was identified that most likely belonged to the Corallicolidae COR2 group (Fig. 6). These ASVs (ASV_16, ASV_19, ASV_22) have a low relative abundance within *P. clavata* but are found across most samples (Fig. 6). This group is most closely related to MN022429_Corallicolidae_COR2_sp. (95.84% Similarity to ASV_16), a sequence recovered from the deep-sea Antipatharian *Leiopathes glaberrima*, also from the Gulf of Mexico. Lastly, a third clade of prevalent corallicolids was identified within *P. clavata* that shows close relation to JX943883_Corallicolidae_COR1_sp. (93.04% Similarity to ASV_105), a sequence found in the stony coral *Porites astreoides,* from the Bahamas.

**Figure 6.**
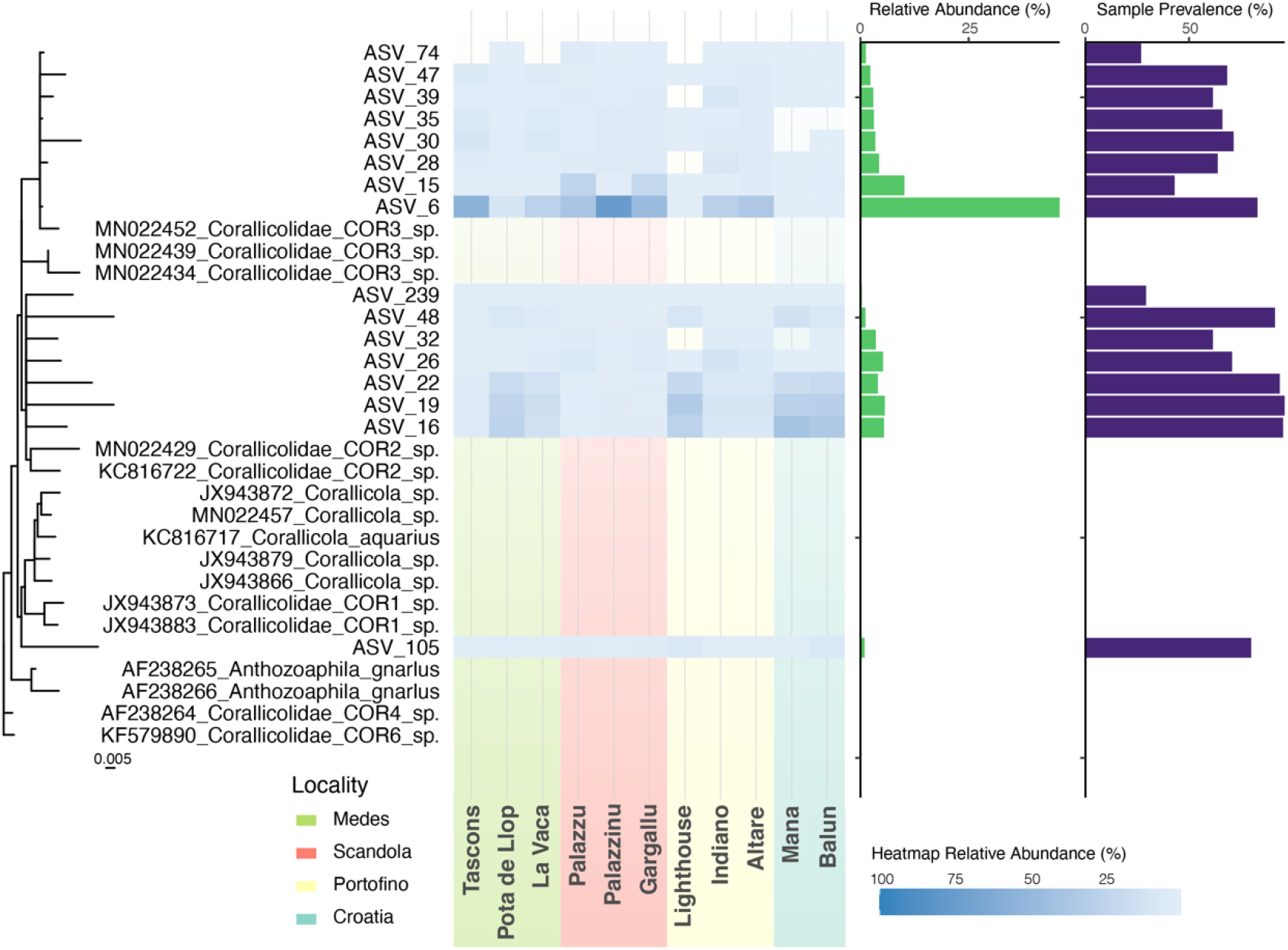
RAxML maximum-likelihood tree of corallicolids. Primary corallicolid ASVs recovered from *P. clavata* placed within the tree structure using EPA. Relative abundance of each ASV at each respective population is shown in the heatmap. Populations are ordered from west to east, with major localities color coded. Relative abundance (green) and sample prevalence (purple) of the ASV across the entire dataset is shown in the barplots to the right.

Examination of the Dinoflagellate ASVs revealed distinct clades present within *P. clavata*. *Dino-Group-I Clade 1* ASVs were most prevalent across the dataset with ASV_13 being the most abundant across all localities except Scandola, where ASV_12 is most prevalent (See Supplementary Figure 5, Additional File 7). The closest blast results for these sequences were to uncultured microeukaryotes isolated from intertidal mudflat sediments and ocean surface water from the East and South China Seas, respectively [26, 27] (100% Similarity). One *Symbiodiniaceae* ASV (ASV_43) was identified in some of the populations (Tascons, La Vaca, Palazzinu, Lighthouse, Altare, Mana & Balun) despite *P. clavata* being azooxanthellate (See Supplementary Figure 5, Additional File 7). The ASV from this study corresponds to *Symbiodinium Temperate Clade A* which has been identified in Mediterranean corals before [28]. It’s closest blast result from NCBI was to a *Symbiodinium sp. Temperate Clade A* sequence (99.21% Similarity) recovered from the stony coral *Mussismilia hispida* [29].

The ciliate ASVs within the dataset revealed three primary clades found variably abundant across the surveyed distribution of *P. clavata*. ASVs corresponding to *Holostichia diademata* (Order: Hypotrichia) (ASV_8) and *Ephelota mammillata* (Order: Suctoria) (ASV_7) were the most abundant and prevalent across the dataset overall (See Supplementary Figure 6, Additional File 8). Interestingly, BLAST results for ASV_8, shows 100% similarity to *Holosticha pullaster* (Order: Hypotrichia) isolated from a freshwater pond in the USA [30]. ASV_7 showed high similarity to *E. mammillata* recovered from *Tubularia* hydroids (98.51% Similarity) [31]. *Licnophora macfarlandi* (Order: Licnophoria), was found across most populations, with its highest abundance being in Mana, Croatia (See Supplementary Figure 6, Additional File 8). Blast results show that these sequences were highly similar to *L. macfarlandi* (92.66% Similarity to ASV_3) recovered from the fluid in the respiratory trees of the giant California sea cucumber *Parastichopus californicus* [32].

### Co-occurrence between members of the microbiome

A microbial co-occurrence network was constructed using SPARCC to analyze the positive correlations between the major microbial families of the *P. clavata* holobiont. *Corallicolidae* & *Dino-Group I Clade 1* had the strongest significant co-association between all the major taxa (Fig. 7). This co-association is of note when you consider these two group’s antagonistic relative abundance patterns across the thermal susceptibility groups (Fig. 4). The three ciliate families also showed some significant positive associations with the corallicolids and *Dino-Group I Clade 1. Endozoicomonas*, *Pseudoalteromonadaceae*, *Vibrionaceaee*, and *Flavobacteriaceae* were the primary bacterial families that showed significant co-occurrence with the different microeukaryotic members of the *P. clavata* holobiont (Fig. 7).

**Figure 7.**
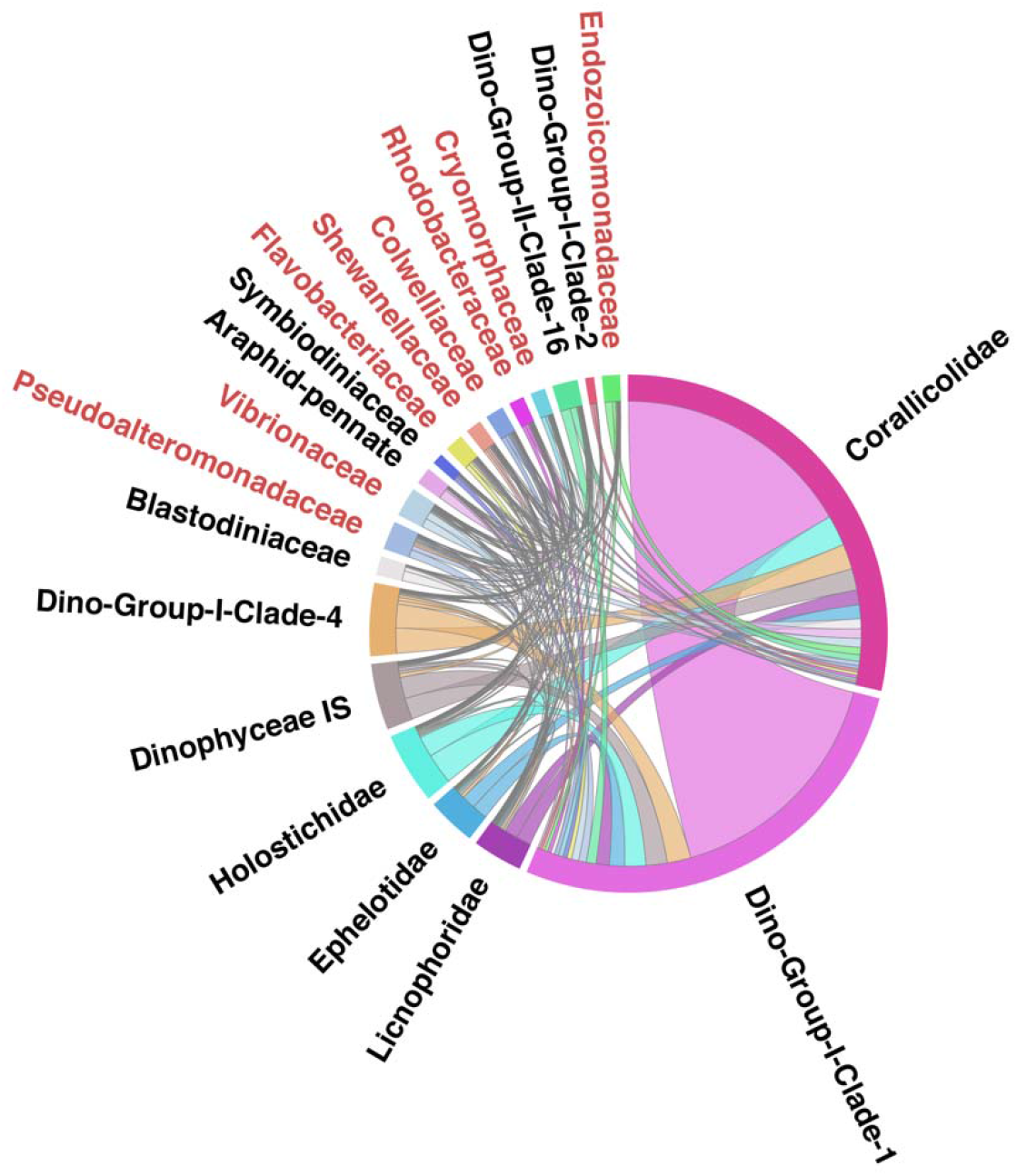
Chord plot of *P. clavata* microbiome. Chord plot of significant positive co-association patterns between the top microbial families across both the 16S and 18S rRNA gene amplicon datasets for *P. clavata*. Interactions were constructed using SPARCC and SpiecEasi with a p-value threshold of 0.05 for significance. Self-interactions have been removed. Eukaryotic families are shown in black text, while Prokaryotic families are shown in red text. IS is short for Incertae sedis, meaning there is uncertainty in the taxonomic position within the broader taxonomic group.

## DISCUSSION

The bacterial community of *P. clavata* showed significant differences in its make-up from location to location, most likely attributed to ASV-level diversity within the *Endozoicomonas* which dominated the bacteriome of each population (Figs. 1, 2, & 4). This is consistent with previous work into the bacterial component of the *P. clavata* holobiont [18, 33]. Additionally, the prevalence of unidentified bacterial ASVs in the dataset points to the underrepresentation of octocoral-associated bacteria in the SILVA reference database. These results exemplify the need to identify more bacteria associated with a healthy symbiotic state in coral holobionts.

The microeukaryotic community of *P. clavata* is dominated by three major groups of alveolates: *Apicomplexa* (mostly corallicolids), *Dinoflagellata* (mostly *Syndiniales*), and *Ciliophora*. This community shows variability in its taxonomic make-up at the order level from location to location, while also showing significant differences in beta-diversity at the ASV level (Figs. 1 & 3). Each of the alveolates found in *P. clavata* showed locality-specific differences, highlighting the specificity and potential co-adaptations of these associations in coral holobionts (Fig. 6, and see Supplementary Figures 5 & 6, Additional Files 7 & 8). Despite being within a clade of obligate parasites, corallicolids are general anthozoan associates found world-wide in healthy corals, even at depths as deep as 1400 m, and in some cases can be as abundant as *Symbiodiniaceae* [34, 35]. They tend to be the most abundant and prevalent microeukaryotic coral symbiont in azooxanthellate corals. The ones recovered from this study showed close relation to those found within deep-sea cnidarians, including octocorals and antipatharians, of the Gulf of Mexico from Vohsen *et al.* [36] (Fig. 6). A third clade of corallicolids were closely related to one recovered from the shallow water Caribbean coral *Porites astreoides* [37]. This diversity of corallicolids present within *P. clavata,* alone, hints at the potential global diversity of corallicolids in other anthozoan hosts.

*Syndiniales*, which have been found in high prevalence in reef environments before, also constituted a major portion of the eukaryotic microbiome. They are an Order of parasites found at the base of the dinoflagellates which can infect a variety of marine macro- and microscopic organisms [38]. *Syndiniales* from *Dino-Group I* are one of the main microeukaryotic taxa found in the hard coral *Pocillopora damicornis* [39], however this group is also ubiquitous in global plankton communities [40]. Additionally, *Dino-Group 1 Syndiniales* with identical 18S rRNA V4 sequences may associate with different hosts, suggesting a generalist host specificity for this clade or the need for a different marker gene to decipher its true diversity [41]. *Syndiniales* have the potential to disrupt nutrient cycling [42], and thus could likely play a role in (de)stabilizing symbiotic relationships, which rely on nutrient dynamics, in a tight-knit symbiotic system such as a coral holobiont. Their differential abundance within *P. clavata* across different thermal susceptibility categories highlights the necessity of studying the role of parasitism in the nutritional dynamics of marine holobionts, especially in the face of stressors. Hudson et al. 2006 noted that a healthy system may be one that is rich in parasitic species due to the impact of parasite-mediated effects on population dynamics, interspecific competition, and energy flow [43]. Applying this perspective to coral holobionts may help us better understand the higher thermal tolerance of *P. clavata* rich in potential parasites such as the corallicolids or *Syndiniales*. The high abundance of these microbes within the *P. clavata* holobiont may contribute to a healthy and resilient system by influencing the microbial community dynamics when confronted with heat-stress.

Within the ciliate lineages recovered from *P. clavata*, three genera (*Licnophora*, *Holosticha*, & *Ephelota*) showed variable abundances across the surveyed distribution of the coral host. *L. macfarlandi*, has been found as an endosymbiont in the respiratory trees of holothuroids, but also in association with corals, likely as a secondary invader hunting other microbes [44, 45]. It is a noted predator of protozoans and is speculated to be attracted to other ciliate species found at the interface of coral lesions [45]. *L. macfarlandi* within *P. damicornis* also exhibited geographic variability in abundance [39], which may depend upon the presence of other ciliates on which it preys. Another ciliate, *H. diademata,* has also been found within corals often crawling slowly on the coral surface [45]. It is a noted bacterivore so it may feed upon the bacteria of the coral, exerting top-down control upon that portion of the microbiome, and therefore influencing the cycling of certain nutrients in the holobiont. The presence of bacterivores, such as ciliates, may increase microbial activity and turnover within the ecosystem [46]. While the previous two ciliates are predominately associated with diseased corals [47], the third ciliate genera, *Ephelota,* has never previously been described in corals; however, suctorian ciliates are typically commensal and have previously been recovered as epibionts of crustaceans, mollusks, fishes, and even other protists [48]. Thus, it may be using the stability and structure provided by the *P. clavata* host to stabilize itself in the water column and capture prey more readily.

This study identified the key microbes associated with *P. clavata,* an important habitat-forming species impacted by MMEs, and provides evidence of the microbiome’s potential role in the thermal susceptibility of the host. In *P. clavata*, the overwhelming abundance of *Endozoicomonas* masked any major differences in bacterial composition. On the other hand, the microeukaryotic community has distinct taxonomic composition associated with different degrees of thermal tolerance. *Dino-Group I Clade 1* was significantly enriched in thermally tolerant corals, while corallicolids were significantly enriched in thermally sensitive corals. These results may explain inter-individual variation in thermotolerance within *P. clavata* populations observed by Gómez-Gras et al. 2022 [15]. How the relative abundance of these apicomplexans confers thermal susceptibility to the coral host is not yet understood. There’s the possibility that the symbiosis between corallicolids and their coral host shifts from an apparent commensalism towards parasitism as added stressors, such as heat, breakdown the relationship. The context-dependency of this relationship can explain the abundance of corallicolids in apparently healthy coral. Briefly, the coral host may have evolved resistance to corallicolids at a significant fitness cost leading to a state of commensalism without the presence of addition stressors; however, heat-stress may cause these corallicolid-tolerant corals to exhibit higher mortality as the apicomplexans shift towards a parasitc relationship with the host, similar to the apparent commensalism of parasites idea proposed by Miller et al. 2006 [49]. This potential interaction and the widespread distribution of the corallicolids would have significant implications when interpreting threats to corals in the context of warming ocean temperatures.

Showing strong co-association with the corallicolids were the *Syndiniales*, specifically those belonging to *Dino-Group I Clade 1* (Fig. 7). These parasites are known to infect a variety of multicellular and microbial hosts and have previously been recovered from coral reef sediments, plankton, and corals [50]. Members include microeukaryotic parasites such as *Duboscquella sp.* which is known to parasitize ciliates [51]. This opens the possibility that they may be infecting either the coral host, it’s microeukaryotic members, or both. While significant positive co-association between *Dino-Group I* and the ciliates recovered from *P. clavata* was apparent in the network analysis, the association between *Dino-Group I* and the corallicolids was most prevalent (Fig. 7). While this can simply be the result of fluctuations in two of the most highly abundant microeukaryotes found in the dataset, it may also represent an interesting case of parasitism of a coral endosymbiont by another member of the microeukaryotic community. Its association with the corallicolids hints at a possible positive role of this parasite preventing the overabundance of corallicolid growth in the coral holobiont which may be associated with decreased thermal tolerance. *Syndiniales* have been proposed to have the capacity to control the sizes of their host populations since a singular infection can produce hundreds of dinospores to further infect new hosts [52]. The vast diversity of known microeukaryotic hosts of *Syndiniales* combined with the larger cell size of corallicolids (∼12 um in length [22]) supports the possibility of this relationship. Possible explanations for this interaction could be top-down control of the corallicolid population by the *Syndiniales*, niche competition, easier co-infection of an already infected host, or excess dinospore production by *Syndiniales*. In either case, the relationship between *Dino-Group I*, the corallicolids and the coral host needs to be investigated further to decipher the ecological relevance of this potentially pivotal interaction within the coral holobiont.

## Conclusions

We have shown here that a deeper characterization of the, often overlooked, microeukaryotic portion of the coral microbiome can have major implications on coral health in a warming ocean. The recently described corallicolids are significantly correlated with thermally susceptible *P. clavata*, yet their role within coral holobionts globally has yet to be uncovered. Along with other genetic, environmental, and physiological mechanisms the microbiome, especially the microeukaryotic portion, should be considered a vital component when assessing coral thermal response. Future research should focus on identifying the mechanisms by which these microbes may confer thermal resiliency to individual corals and how this knowledge can best be used to manage fragile coral ecosystems.

## Declarations

### Ethics approval and consent to participate

Not applicable.

### Consent for publication

Not applicable.

### Availability of data and materials

The raw sequencing data for this project has been deposited on NCBI (BioProject: PRJNA928446). All bioinformatic scripts used in this study can be found at https://github.com/Abonacolta/Bonacolta_2022_Pcla.

## Authors’ Contributions

JdC conceptualized the study. JdC, DG, JBL, PL, JG collected the samples and performed the experiment. JdC and JM prepped the samples for sequencing. AMB conducted the formal analysis, generated the figures, and wrote the manuscript. JdC, DG, JBL, PL, JG, and R.M provided the laboratory resources for the experiment. All authors read and approved the final manuscript.

## Acknowledgments

We appreciate the contributions of the following scientists in helping with our sampling effort: Bensoussan, N, Cerrano C, Ferretti E, Kipson S, Bakran-Petricioli T, Serrao EA, Paulo D, Boavida J, Montero-Serra I, López-Sanz A, Milanese M, Linares C.

## Additional Information

**Supplementary Figure 1.**
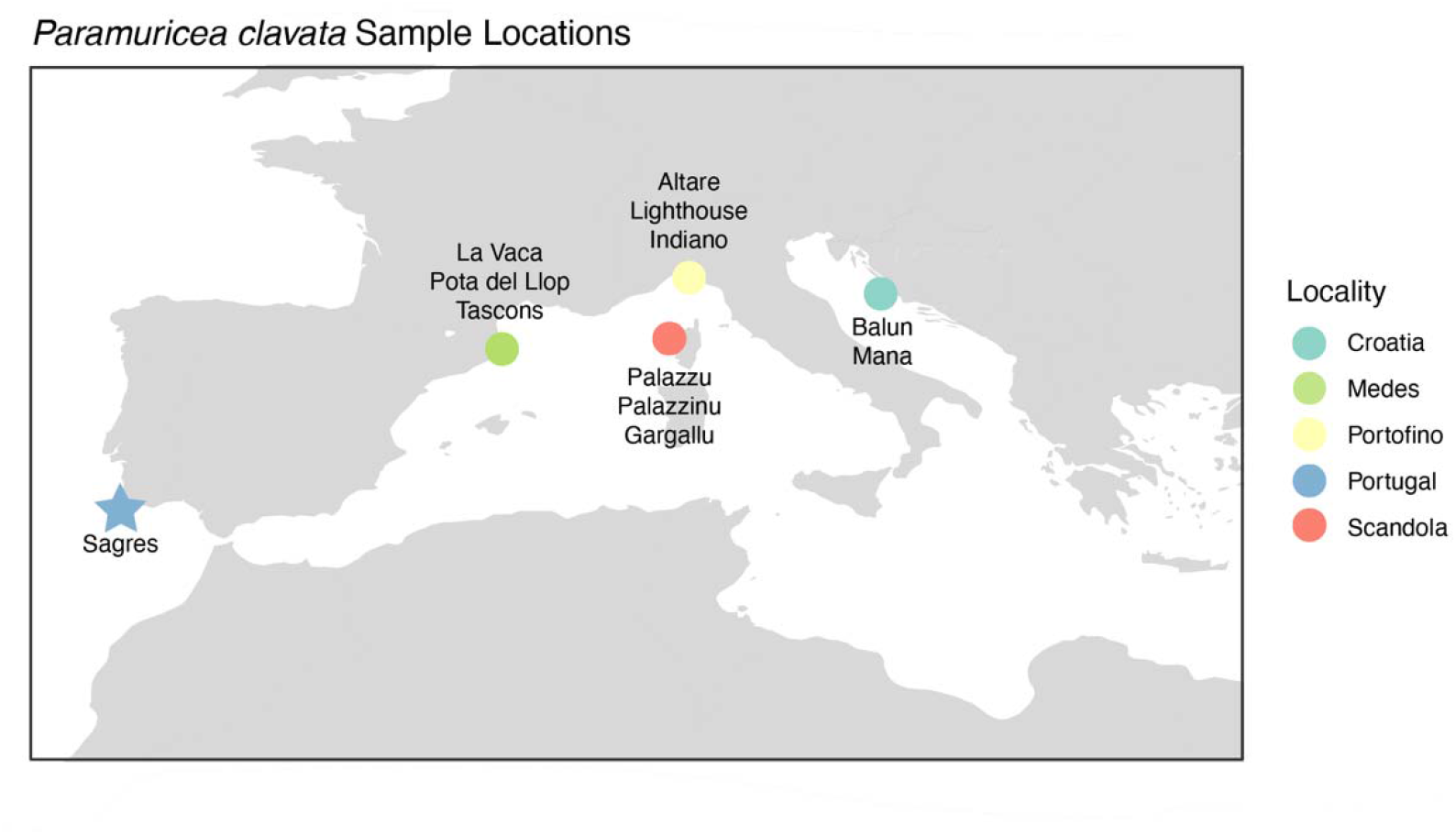
Additional File 1. Map showing locations of sampled localities of *Paramuricea clavata*. Sagres is denoted by a star as this population was later determined to be a different species.

**Supplementary Table 1 Additional File 2, Sample Metadata.** Excel format (.xls). Information on each coral specimen analyzed in this study including thermal-tolerance categories, necrosis percentages, origin name and location, as well as environmental characteristics of population like depth and temperature at time of sampling.

**Supplementary Table 2 Additional File 3, Thermal Categories System.** Excel format (.xls). Information on how each coral’s mortality was assessed and categorized at days 14 and 21 of the heat-stress experiment. This table also shows how these two measurements were interpreted for the final thermal tolerance category for the coral fragment.

**Supplementary Figure 2.**
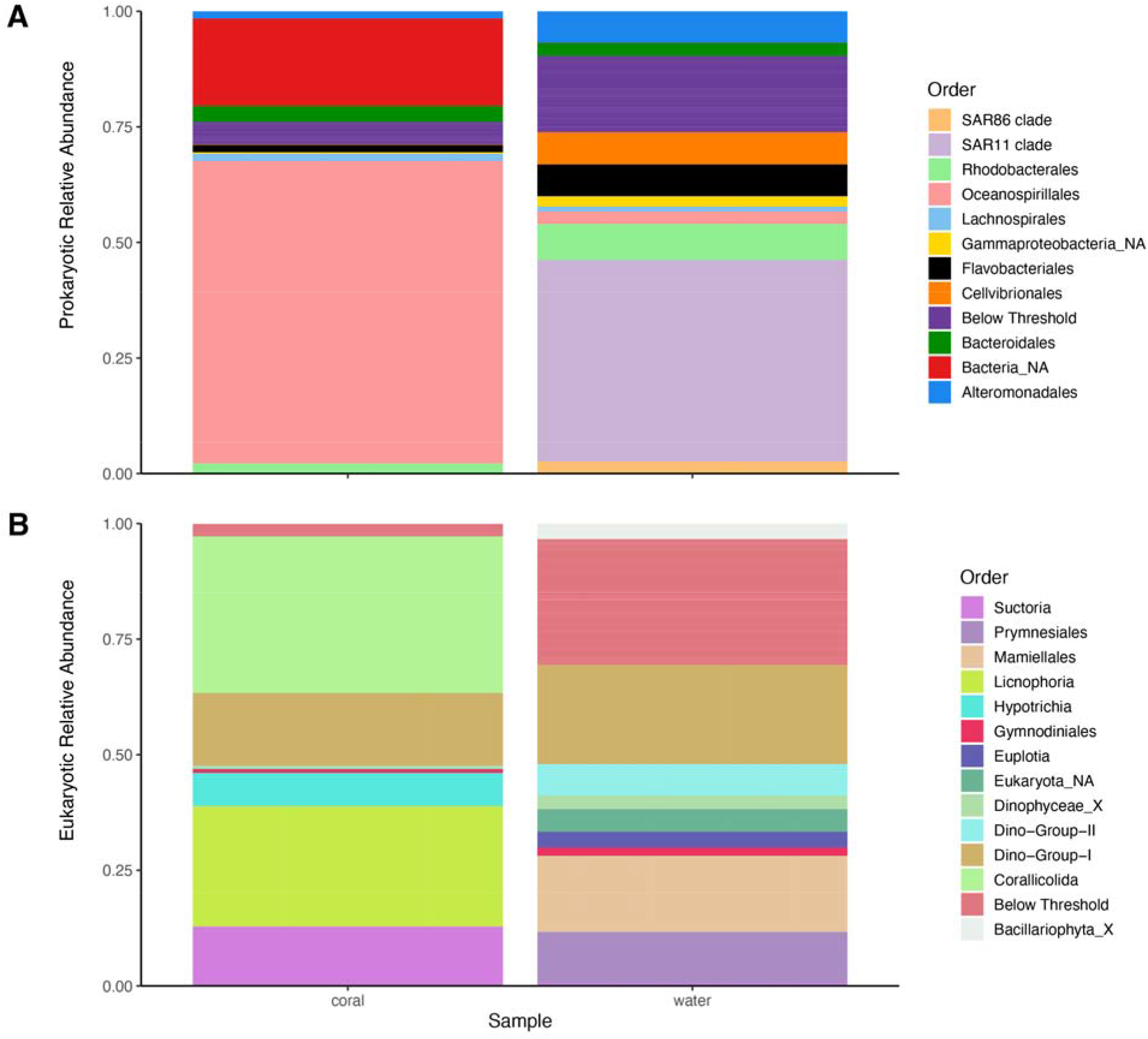
Additional File 4. Stacked bar plots comparing the microbial order composition of *P. clavata*. The prokaryotic (A) and eukaryotic (B) communities of the coral vs. the water it was sampled from is depicted. Only those taxa with a median relative abundance above 1% are shown, the rest are listed as “Below Threshold”.

**Supplementary Figure 3.**
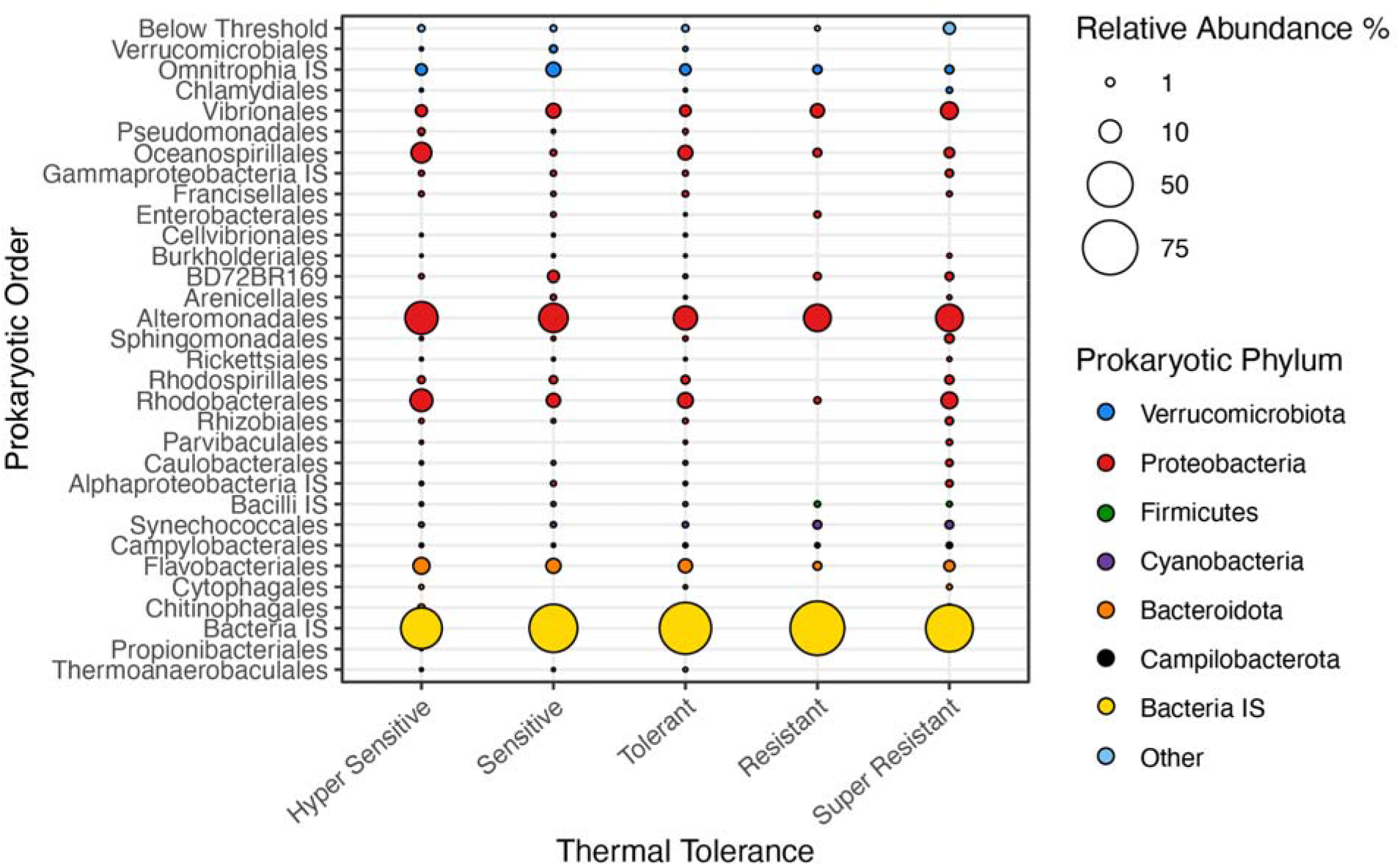
Additional File 5. Relative abundance of bacterial orders within *P. clavata* across thermal tolerance categories with the highly abundant *Endozoicomonas* removed. Hyper Sensitive corals were those which showed 100% necrosis after the heat-stress experiment. Super Resistant corals showed 0% necrosis at the end of heat-stress. Microbial taxa are ordered taxonomically. Only those taxa with a median relative abundance above 1% are shown, the rest are listed as “other”. IS is short for Incertae sedis, meaning there is uncertainty in the taxonomic position within the broader taxonomic group. Relative abundance bubbles correspond to within domain relative abundance. The high abundance of Bacteria IS can be contributed to poor representation of octocoral bacteria in the SILVA reference database.

**Supplementary Figure 4.**
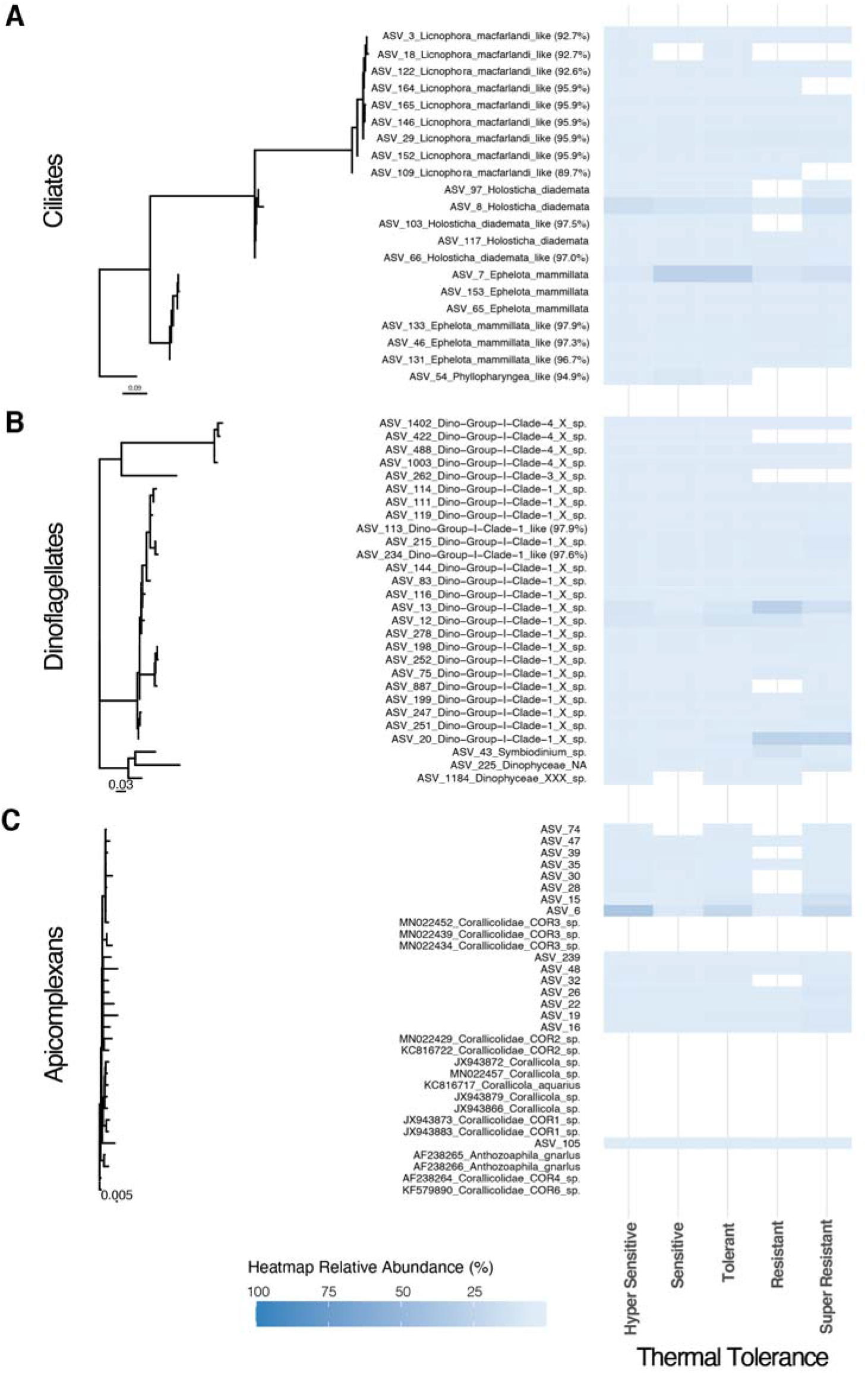
Additional File 6. Maximum-likelihood tree of top alveolate ASVs across the thermal tolerance categories of *P. clavata.* Heatmap abundance account for the relative abundance of each specific ASV across the alveolates shown.

**Supplementary Figure 5.**
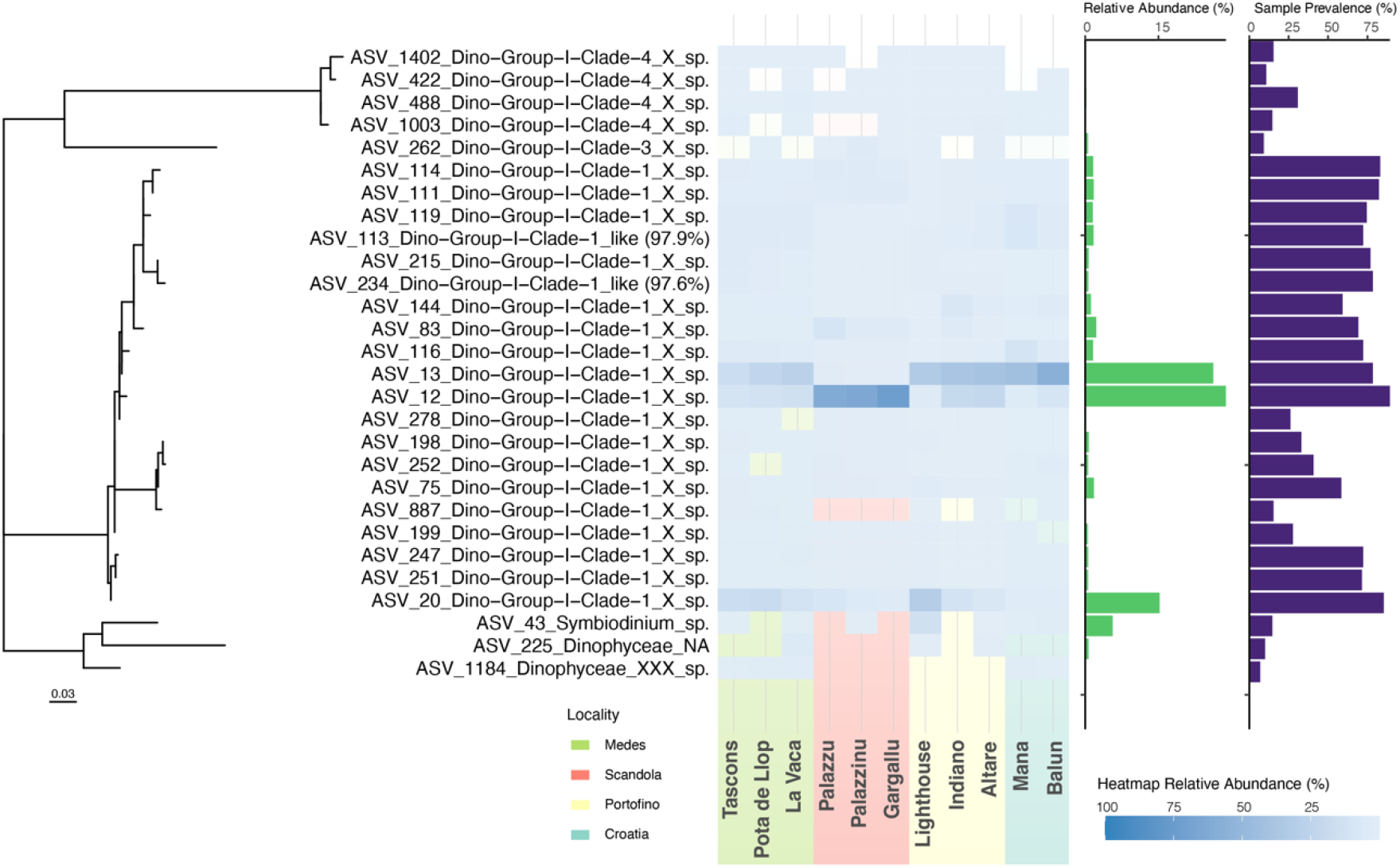
Additional File 7. Maximum-likelihood tree of top dinoflagellate ASVs across the sampled populations of *P. clavata.* Populations are ordered from west to east, with major geographic localities color coded. ASV taxonomy is assigned to species level if sequences share above 98% similarity with their reference. ASV abundance is also shown across the whole data set (green bars) as well the ASV prevalence across all the samples (purple bars).

**Supplementary Figure 6.**
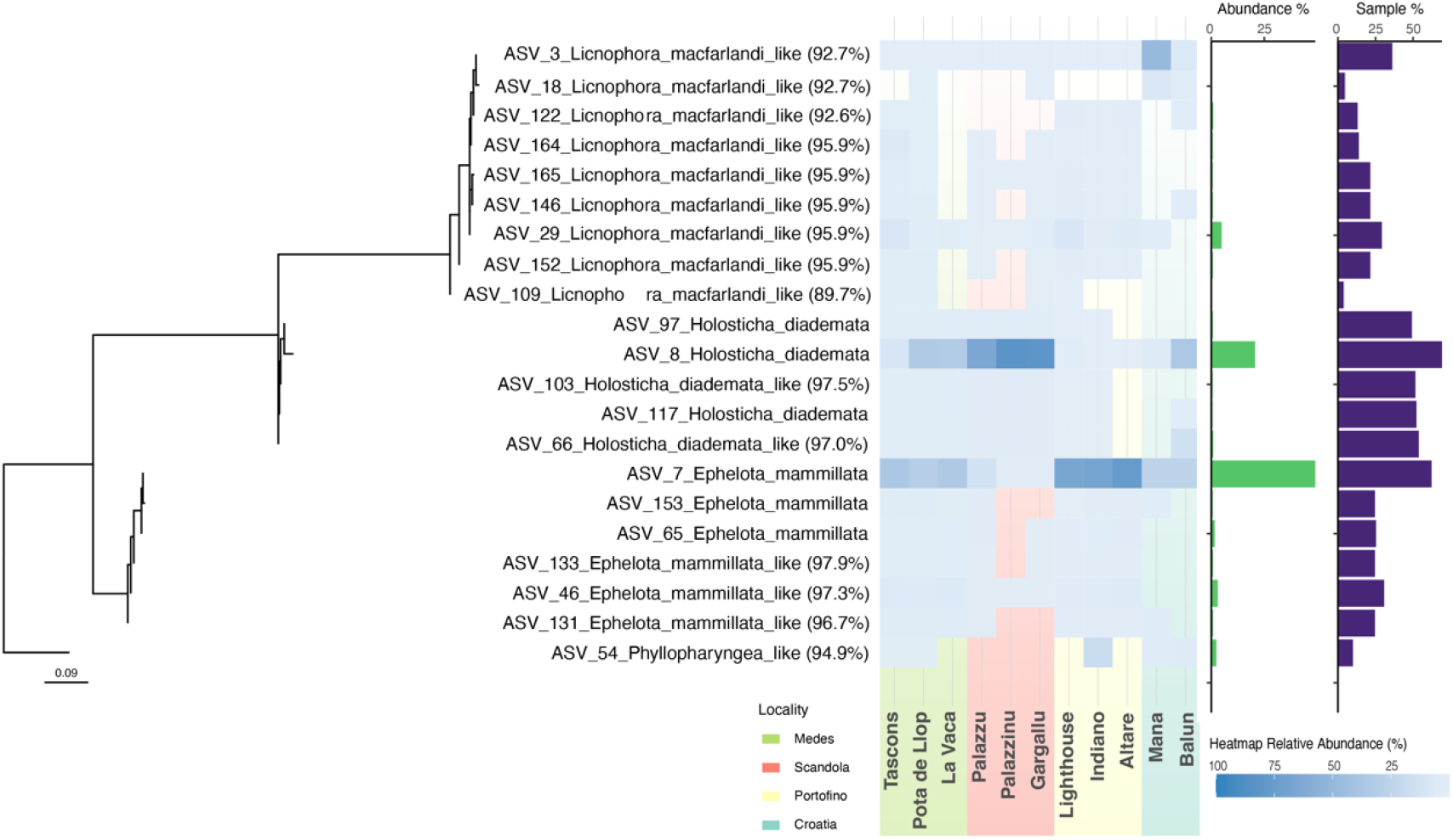
Additional File 8. Maximum-likelihood tree of top ciliate ASVs across the sampled populations of *P. clavata.* Populations are ordered from west to east, with major geographic localities color coded [Medes = Green, Scandola = Red, Portofino = Yellow, Croatia = Cyan]. ASV taxonomy is assigned to species level if sequences share above 98% similarity with their reference. ASV abundance is also shown across the whole data set (green bars) as well the ASV prevalence across all the samples (purple bars).

